# Expression and T cell Regulatory Action of the PD-1 Immune Checkpoint in the Ovary and Fallopian Tube

**DOI:** 10.1101/2020.06.06.138123

**Authors:** Joshua Johnson, Peter Ka Sam, Rengasamy Asokan, Evelyn Llerena Cari, Elise S. Bales, Thanh-Ha Luu, Lauren Perez, Amanda N. Kallen, Liesl Nel-Themaat, Alex J. Polotsky, Miriam D. Post, David J. Orlicky, Kimberly R. Jordan, Benjamin G. Bitler

**Affiliations:** University of Colorado Anschutz Medical Campus, Department of Obstetrics and Gynecology, Division of Reproductive Sciences, Building RC2, Room P15 3103, Mail Stop 8613, Aurora, Colorado 80045; University of Colorado Anschutz Medical Campus, Department of Obstetrics and Gynecology, Division of Reproductive Endocrinology and Infertility, 12631 East 17th Avenue, Room 4409, B198-3 Aurora, Colorado 80045; University of Colorado-Boulder; University of California-Los Angeles; Yale School of Medicine, Department of Obstetrics & Gynecology; University of Colorado Anschutz Medical Campus, Department of Pathology, Mailstop F768, 12605 East 16th Avenue, Aurora, Colorado 80045; University of Colorado Anschutz Medical Campus, Department of Immunology and Microbiology, Human Immunology and Immunotherapy Initiative, Human Immune Monitoring Shared Resource, RC1-North, 8113, Aurora, Colorado 80045

**Keywords:** folliculogenesis, immune checkpoint, immunotherapy, ovarian cancer, ovary, menopause, PD-1

## Abstract

The <u>P</u>rogrammed Cell <u>D</u>eath Protein-1 (PD-1/PDCD-1/CD279) checkpoint has powerful immunomodulatory action, including in the context of cancer. PD-1 receptor activation by its ligands (PD-L1/2) is associated with downregulated immune response, and tumor cells can avoid surveillance *via* PD-1 and/or ligand expression. While receptor expression is largely limited to lymphoid, myeloid, and tumor cells, we show that membrane bound and soluble variants of PD-1 and ligands are also expressed by permanent constituent cell types of the human ovary and fallopian tube, including granulosa cells and oocytes. PD-1 and soluble ligands were highly enriched in exosome fractions in human follicular fluid at bioactive levels that can control T cell PD-1 activation. PD-1 checkpoint signaling may be involved in physiological ovarian functions including follicle, and ultimately, germline and embryo immune-privilege.

## Introduction

There is a critical need to minimize autoimmune responses to oocytes or to other cells that might compromise fertility^1,2^. The clinical condition termed autoimmune oophoritis is associated with poorer fertility outcomes. The condition manifests in immune cell infiltration of the ovaries, which can take on a cystic appearance^3,4^. Autoimmune oophoritis can include the presence of anti-ovarian antibodies, and often occurs when another definitive autoimmune condition (e.g., lupus, autoimmune polyglandular syndrome, etc.) has been diagnosed. Very recently, a group has shown that there are decreased numbers of effector T regulatory (Treg) cells and increased CD4+ CD69+ T cells in the peripheral blood of women afflicted with primary ovarian insufficiency^5^, a condition characterized by accelerated ovarian aging early loss of ovarian function.

The PD-1 immune checkpoint^6–8^ controls immune cell (T^9^, B^10^, and dendritic^11^ cell) activation, favoring identification of interacting cells as “self,” and thus immunosuppression, when induced. With rare exceptions^12^, the PD-1 receptor has only been detected in myeloid and lymphoid cells, and in tumor cells that co-opt the pathway to evade surveillance and elimination^8,13^. Transformed cells can downregulate anticancer immune responses by expressing PD-1^14^, ligands PD-L1^15–17^ or PD-L2^18^, or soluble variants of these proteins^19–22^. We hypothesized that if present, local PD-1 signaling in the ovary and fallopian tube could supppress autoimmunity to native cells including oocytes (within the ovary) and embryos (trafficking within the tubal lumen). PD-1 signaling in the ovary and fallopian tube might also then be involved in the development of early stages of epithelial ovarian cancer (EOC).

PD-1 pathway expression in EOC cells has been shown to correspond to disease stage and patient survival in this manner^23–26^, but very little information about expression in normal ovarian or tubal cell types has been made available. Notably, EOC is considered an immunogenic disease^27^, but the response rate to checkpoint inhibitor immunotherapies has been unexpectedly low^28^.

The most common histotype of EOC is high grade serous ovarian cancer (HGSOC). The originating cells of HGSOC are increasingly thought to be transformed fallopian tube epithelial (FTE) cells^29–35^. Early transformation events can result in <u>s</u>erous tubal intraepithelial lesions (STILs), which are characterized by a “p53 signature” in morphologically unremarkable FTE cells, visible on immunohistochemical staining as overexpression in 12 or more consecutive non-ciliated cells. Mutations in the tumor suppressor can result in either nuclear stabilization of the mutant p53 variant, or, in loss of p53 expression^36^. STILs can then progress to another stereotypical stage, the <u>s</u>erous tubal intraepithelial <u>c</u>arcinoma (STIC)^30,37,38^. STIC cells detach from the tubal lumen and engraft at distant sites within the peritoneum, including the surface of the ovary at ovulation site(s)^39,40^.

During a preliminary evaluation of Pd-1 pathway protein expression in immune cells within the mouse ovary, we detected broad expression of the receptor and ligands in the organ, in both somatic cells and oocytes. There was also an existing report in a publicly-available large-scale gene expression screen (LifeMap Discovery^®^) that showed that PD-1 pathway transcripts are detectable in human cumulus granulosa cells (HCGC). Because of potential relevance of these initial data to ovarian function and ovarian cancer development, a more extensive analysis of the PD-1 checkpoint in the human ovary and fallopian tube was warranted. We tested whether the PD-1 checkpoint acts within the human ovary and fallopian tube in ways suggestive of physiological function, including testing whether soluble PD-1 or ligand proteins are present in human follicular fluid (HFF) at levels capable of regulating immune function.

## Results

In an immunohistochemical study of both pre- and postmenopausal human ovarian tissue (Fig. 1), we found that PD-1, PD-L1, and PD-L2 are are widely expressed, and this was conserved in the mouse ovary (Fig. S1, human tonsil positive controls shown in Fig. S2). In human premenopausal specimens (n=8 unique patient samples evaluated), PD-1 was detectable in the oocytes of non-growing (including primordial stage) and growing follicles (Fig. 1a,b), but unlike the mouse (Fig. S1), some oocytes appeared negative (examples in 1b). Like the mouse, PD-1 was present at low relative levels in granulosa cells of growing follicles (red arrow, 1b,c; see also 4e), and cells of the ovarian cortex (red arrows 1d). The receptor was also detected in what appeared to be immune cells subjacent to the fallopian tube lumen (1e, denoted by *), with epithelial cells of the tubal lumen faintly positive (arrow, 1e’). In addition, cells positive for PD-1 with the appearance of immune cells were detectable throughout the stromal/extrafollicular tissue of each ovary specimen (arrowheads, 1b,d), and their identities were probed by double immunofluorescence staining, and, multispectral imaging (Vectra). Macrophages double-positive for PD-1 and CD68 were detected, and T cells double-positive for PD-1 and CD3 were detected in ovary specimens (Fig. S3).

**Figure 1.**
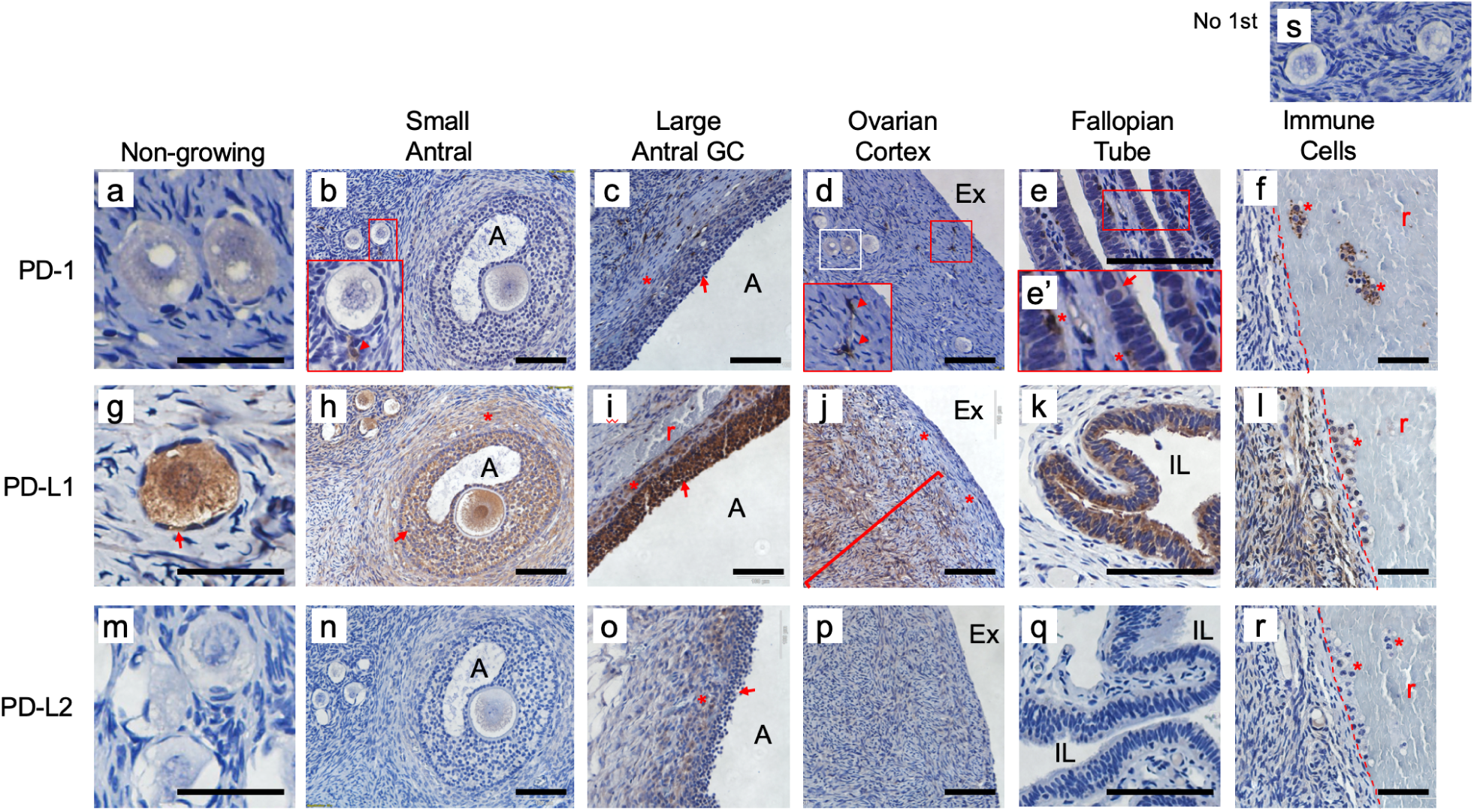
Premenopausal expression of PD-1 receptor and ligands in human ovarian follicles and fallopian tube. Brown stain indicates expression. PD-1 was detected in some oocytes (compare faintly positive examples in panel a with negative examples in b and f; region from white box in panel d is shown in panel a), and granulosa cells of large antral follicles (c, arrow; theca interna denoted by *). PD-1 was present at high levels in relatively rare cells throughout the ovary (red arrowheads in insets, b, d) and the stromal compartment of the fallopian tube (inset e’, *); some of these are likely to be T cells and macrophages (see Figure S2). PD-1 was also detectable at low levels in the luminal epithelium of the fallopian tube (e and e’, arrow). Control circulating immune cell expression of PD-1 is shown in panel f (cells within ovarian blood vessel marked by *; inner endothelial lining marked by dashed red line in panels f, l, and r). PD-L1 was consistently expressed in oocytes (g,h) of all sizes, pregranulosa cells of non-growing follicles (g, arrow) and granulosa cells (h, i, arrows) and the theca interna (h, i, *) of growing follicles. PD-L1 was also broadly expressed in stromal cells of the inner ovarian cortex (j, l – left), but was not detectable in cells of the OSE or in stromal layers immediately below the surface (j, red bracket and *’s; compare to Fig. 2e-g). PD-L1 was also expressed the luminal epithelial cells of the fallopian tube (k), and circulating immune cells (l). PD-L2 was detected only in the granulosa cells (arrow) and theca interna (*) of large antral follicles (compare panel o to m,n,p,q,r). No first antibody control shown in panel s. A – follicular antrum, Ex – extra-ovarian space, IL – intraluminal space of tube, r – red blood cells. Scale bars in panels a,e,g,k,m,q = 100 *µ*m, bars in other panels are 50 *µ*m

The PD-1 activating ligand PD-L1^15–17^ was expressed broadly in the premenopausal ovary and fallopian tube. PD-L1 was detected in all oocytes in all specimens (examples in 1g,h and 4e), in granulosa cells of follicles of all sizes (1g-i, arrows, also 4e), cells of the *theca interna* of growing follicles (1h,i *), cells of the ovarian cortex (1j, compare to 1d and 1p), and FTE cells (1k). PD-L1 expression in the stromal cells of the premenopausal ovarian cortex was consistently found *not* to extend to the ovary surface; instead, PD-L1 was absent from the layers of the ovary closest to surface. PD-L2 was less widely expressed in these cell types compared to PD-L1, with all oocytes and granulosa cells of follicles up to the small antral stages negative (1m,n), and only granulosa cells of larger follicles (1o, arrow; Fig. 4f) and cells of the *theca interna* (1o, *) were consistently positive.

**Figure 2.**
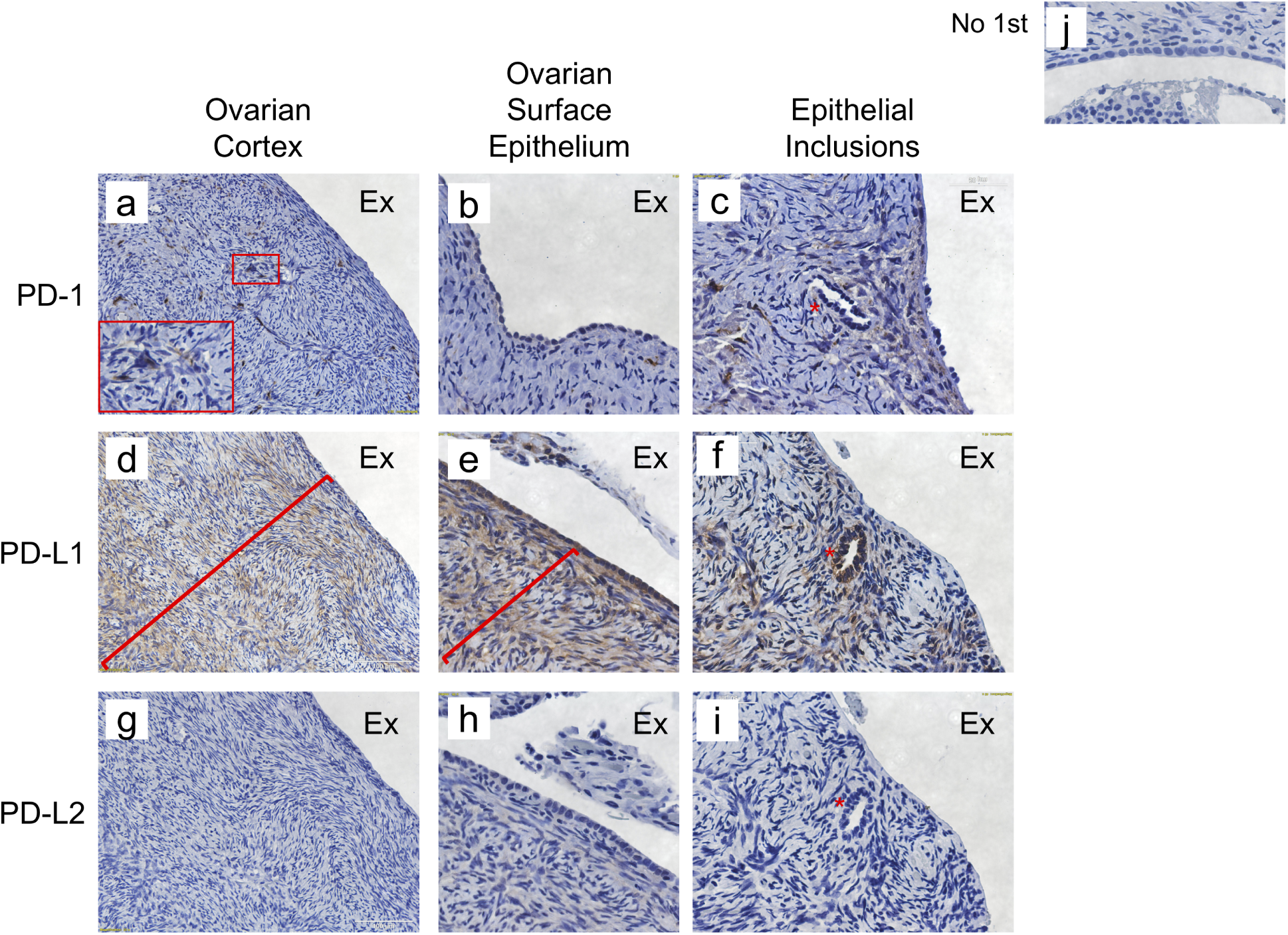
Postmenopausal human ovarian PD-1 pathway expression. As seen with premenopausal specimens, PD-1 expression was highest in sparse cells with the appearance of surveilling immune cells in the ovarian cortex (a, inset), but was low or absent in cells of the OSE (b) and sub-surface epithelial inclusions (c, *). PD-L1 was widely expressed throughout postmenopausal ovarian cortex (d-f), and unlike the premenopausal ovary, was detectable at comparable levels in cell layers at the surface of the ovary (red brackets, panels d,e; compare to Fig. 1j). PD-L1 was also detected in cells of the OSE (d,e) and in cells of epithelial inclusions (example in f, note lack of surface OSE in this area). PD-L2 was mostly absent from postmenopausal specimens (g-i). No first antibody control shown in panel j. Ex – extra-ovarian space.

**Figure 3.**
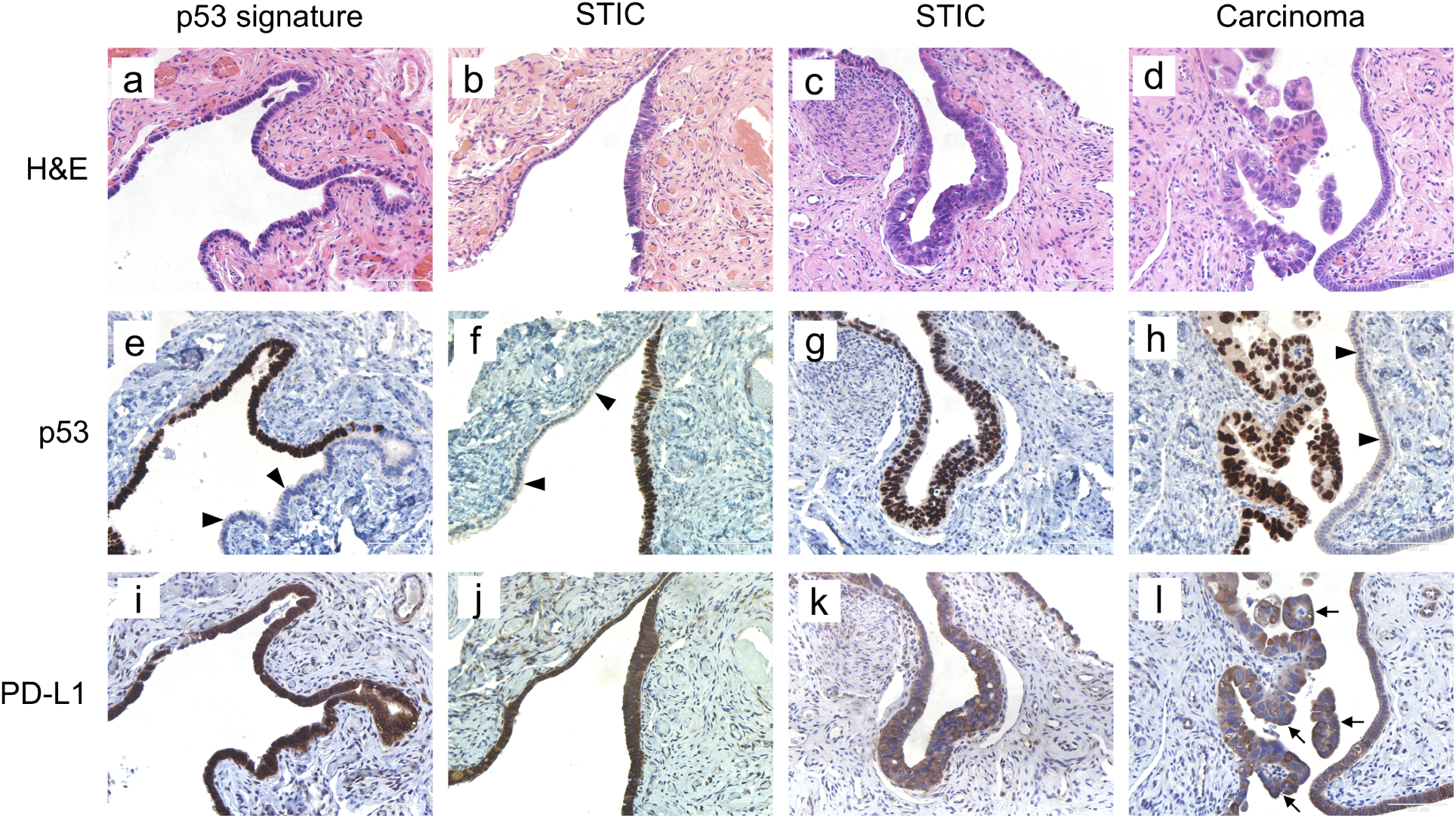
PD-L1 expression in p53-variant tubal epithelial lesions. As seen in the normal fallopian tube (Fig. 1k), PD-L1 is expressed in tubal lesions that have been implicated as precursors of HGSOC. Hematoxylin and eosin stained sections used for clinical/pathological staging are shown in panels a-d, with adjacent serial sections stained for p53 (e-h) and PD-L1 (i-l) below. Both normal (p53 low/absent, arrowheads e,f,h) and p53-overexpressing (brown staining, p53 row of images) FTE cells were positive for PD-L1, in lesions of all types evaluated.

**Figure 4.**
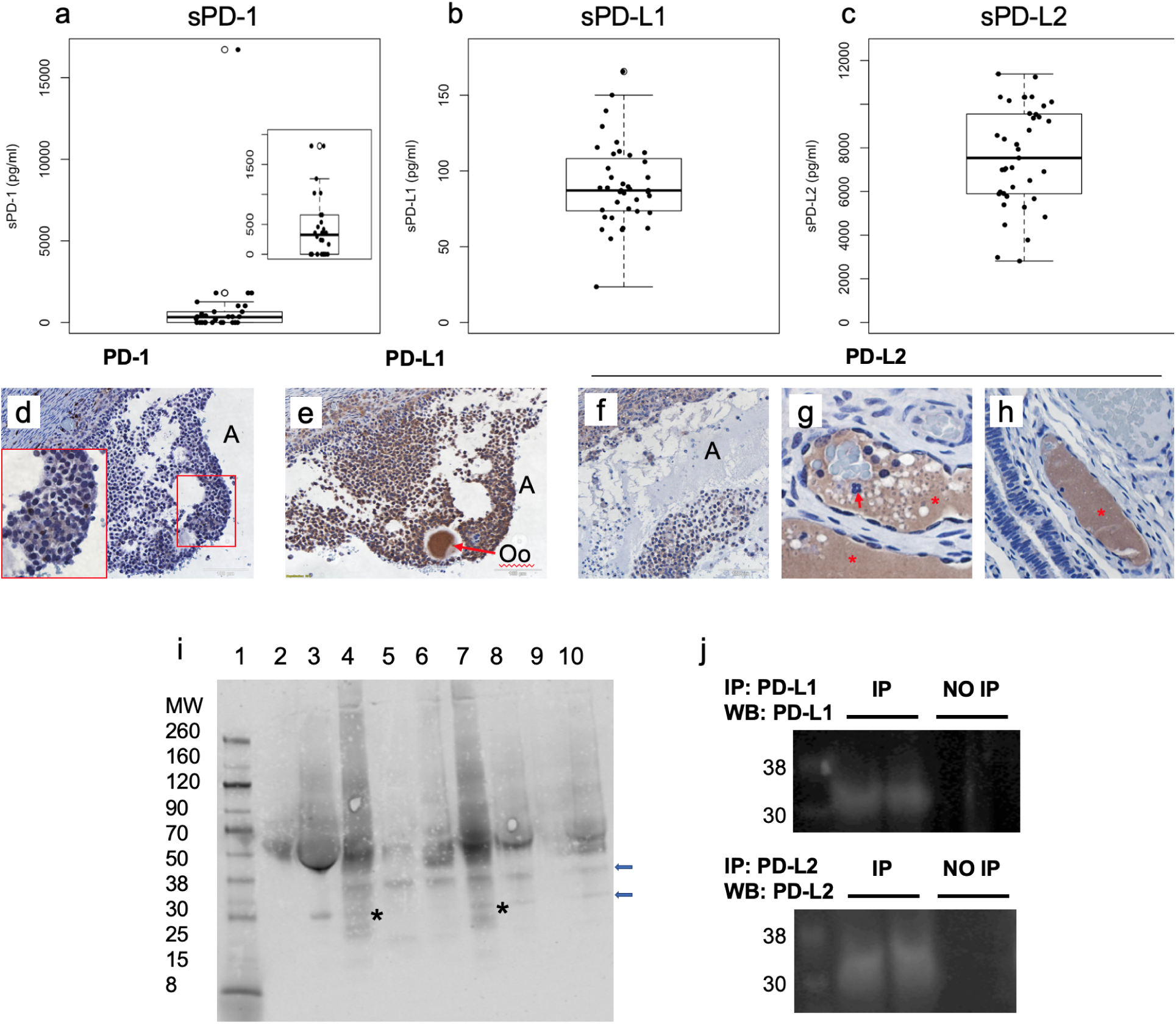
Detection of PD-1 receptor and ligands in human follicular fluid. ELISA assays were used to determine whether sPD and/or PD-1 ligands are present in HFF. PD-1 was detected in 20 of 32 samples measured (a, inset plot shows same data points between 0 and 1750 pg/ml expanded vertically, note single sample with greater than 15,000 pg/ml PD-1), while PD-L1 (b) and PD-L2 (c) were detected in all samples. Subsequent analysis detected full-length and shorter variants of each protein as enriched in HFF exosomal fractions (see Fig. 5). Representative examples of granulosa cells (gc) in large antral follicles, including HCGC are shown in panels d-f for each protein; this expression pattern may represent membrane and soluble forms of the proteins. PD-L2 was also detected consistently in cell-free contents (*) of ovarian (g) and fallopian tube (h) lymphvascular channels. Note that in the example ovarian lymphvascular channel (g), white blood cells are negative for PD-L2 (arrow), while the cell-free vessel contents in the fixed tissue are positive. Also note that the contents of the fallopian tube channel (h, *) are positive for PD-L2 while the contents of the vein at top right are negative; tubal luminal epithelium is at bottom right. (i) Western blot analysis of full-length and soluble PD-1 in HFF and media conditioned by HCGC, soluble form(s) enhanced by immunoprecipitation (IP) indicated by asterisks. Contents of lanes as follows: Lane 1: MW Marker, 2: media conditioned by HCGC (“HCGC media”) for 24 hours, 3: HFF, 4: HCGC media, anti-PD-1 IP, 5: HCGC media, IP no-first antibody control 6: HCGC media, unrelated first antibody control, 7: HFF, anti-PD-1 IP, 8: HFF, IP no-first antibody control, 9: HFF, unrelated first antibody control, 10: HFF, IP no-first antibody control. (j) Western blot detection of ligands PD-L1 and -L2 in HFF. A first immunoprecipitation (IP) step was required to detect the ligands (equal amounts of protein are loaded with and without IP).

Because the risk of HGSOC increases after menopause^41^, we also evaluated ovarian specimens from postmenopausal patients (Fig. 2, representative data from n=5 unique patient samples shown). In the postmenopausal ovary, PD-1 expression was limited to sparse cells with the appearance of tissue resident immune cells within the ovarian cortex (2a, inset), and was faintly detectable in cells of the ovarian surface epithelium (OSE) (2b) or in sub-surface epithelial inclusions (2c,d). PD-L1 was expressed throughout the ovarian cortex (2e,f), including stromal cells near the organ surface and within cells of the OSE (2f). PD-L2 was not detected in any of the same cell types or structures evaluated in postmenopausal specimens (2i-l).

Despite their having been removed from patients for reasons unrelated to ovarian pathology, 4 of the 5 postmenopausal human ovary specimens harbored surface epithelial inclusions (2c,f,i) thought to develop *via* invagination of cells normally at the ovarian surface^42^. These structures were only positive for PD-L1 (2f).

Given the consistent expression of PD-L1 in normal FTE cells, we also asked whether its expression might change when tubal cells become transformed and acquire a “p53 signature^35,36^.” We limited our analysis to lesions exhibiting a p53 *stabilization* signature, and photomicrographs of hematoxylin-eosin staining as well as PD-L1 immunostaining in serial sections of the same lesion were prepared. Representative images of tubal lesions, including a serial section stained with hematoxylin and eosin are shown in Figure 3. Cells overexpressing p53 (3e) in p53 Signature/STIL lesions were found to maintain expression of PD-L1 (3i). FTE cells within STIC lesions were also PD-L1 positive (panels b,f,j and c,g,k), as were all other cases of more advanced tubal lesions (example more advanced tubal lesion exhibiting mucosal involvement by invasive serous carcinoma, d,h,l). Thus expression of PD-L1 ligand was maintained in the tubal epithelium throughout all stages of transformation evaluated.

The expression of PD-1 pathway proteins pre- and post-menopause were suggestive of ovarian and tubal immunomodulation involving PD-1 checkpoint signaling (Discussion, below). How the pathway might act acutely to regulate immune function remained unclear, and we thus moved to the analysis of potential PD-1 pathway bioactivity in live human cells and fluid isolated from human ovaries in the clinic.

HCGC and (human) follicular fluid (HFF) are readily available as discarded material after clinical egg retrieval procedures. Using the collection technique of Jungheim et al.^43,44^ that allows the collection of i) HCGC and ii) undiluted HFF free of cells, we collected a series of cell samples and fluid aliquots from 60 patients. Where possible, samples were collected from the largest periovulatory follicle in the left ovary, and separately, in the right ovary. Analysis of soluble receptor and ligand levels was first performed by ELISA assay. Soluble PD-1 was detectable in only a subset of HFF samples by ELISA. 12 of 32 samples contained zero detectable PD-1, and samples that did contain PD-1 ranged from approximately 200 pg/ml to greater than 16500 pg/ml (Fig. 4a). In contrast, soluble PD-L1 (4b) and PD-L2 (4c) were detected in *all* HFF samples, and their approximate ranges were between 20 and 160 pg/ml and 2800 and 11000 pg/ml, respectively. Detection of soluble PD-1 receptor and ligands in HFF samples corresponded to their detection in the granulosa cells (including HCGC) seen in immunostained specimens that contained large follicles (4d-f, see also Fig. 1c,i,o). Strikingly, the contents of lymphvascular channels also demonstrated high levels of immunoreactivity for PD-L2 (4g,h). Because the PD-L2 signal was uniform throughout acellular lymphvascular contents, extracellular/secreted PD-L2 is further supported.

Full-length PD-1, along with PD-L1 and PD-L2 protein, as well as shorter splice variants were also detected in each of 4 patient HCGC samples and HFF samples by western blot with optional immunoprecipitation (IP; representative data in 4i,j). In the case of sPD-1 IP, we also assessed conditioned media used to culture HCGC cells for 24 hours (4i). Low molecular weight bands corresponding to the correct sizes of sPD-1 splice variants were detected when anti-PD-1 antibody was used for IP, but these bands were not present when no first antibody, or, an unrelated antibody were used (not shown). IP was required for the detection of sPD-L1 and sPD-L2 on western blots (4j).

Hypothesizing that patients might show a consistent HFF soluble PD-1 pathway “signature,” we first tested whether any correlation between protein concentrations occured i) for the targets between samples collected from the left and right ovary (Fig. S4), ii) for levels of the soluble ligands with each other (Fig. S5a), or iii) if any correlation existed between levels and key patient demographic information (Fig. S5b-d). While PD-1 levels were not correlated between samples collected from the largest follicle in the left versus right ovary (S4a) highly significant correlations for both of the soluble ligands were detected (S4b,c). This suggests that the production of the soluble ligands is a tightly regulated process in individuals. In addition, significant positive correlations were noted between patient age and both PD-L1 and -L2, and patient BMI and PD-L1 (but not PD-L2). No correlation was noted between the number of eggs retrieved and either soluble ligand. These data suggest that at least in the context of *in vitro* fertilization (IVF) treatment, the soluble PD-1 checkpoint immunomodulatory milieu can vary between patients and is subject to modification by aging and body composition. Finally here, the detection of bands corresponding to the sizes of full-length PD-1 and ligands in cell-free HFF (Fig. 4) suggested that an alternative mode of production and delivery of full-length proteins to HFF might be occurring. We hypothesized that full-length proteins might be loaded into exosomes as has been reported for PD-L1^45–48^ and secreted into HFF in that fashion. After processing HFF from 7 unique patients into exosome-enriched and exosome-depleted (soluble protein) fractions, we found that full-length receptor and both ligands, as well as shorter species suggestive of splicing were highly enriched in the exosome fractions of each specimen (Fig. 5).

**Figure 5.**
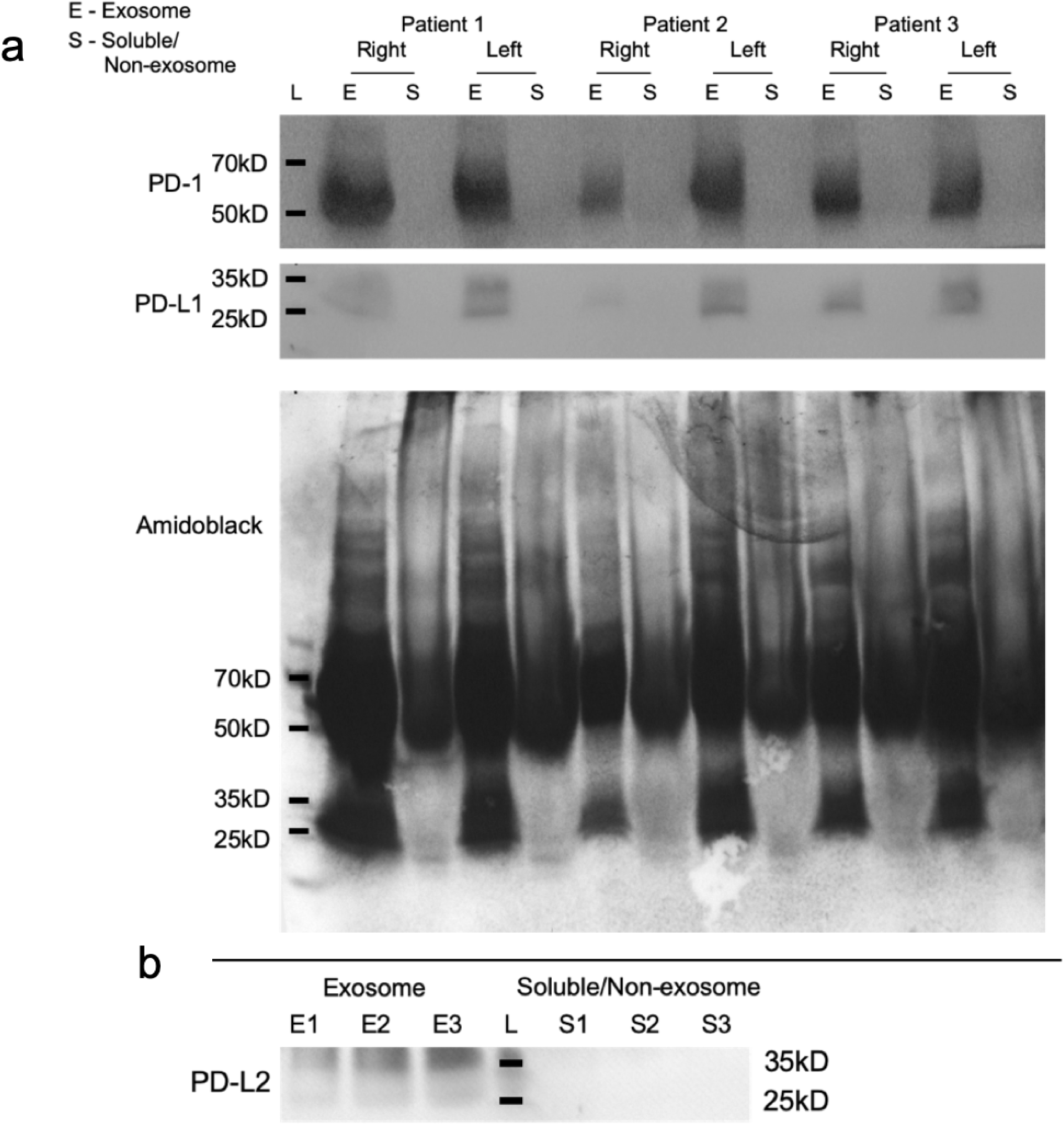
Enrichment of PD-1 and ligands in exosome fraction of human follicular fluid. Partitioning of HFF into exosomal and non-exosome (soluble) fractions revealed that PD-1 and its ligands are highly enriched in the exosome fraction. Panel a shows the detection of PD-1 and PD-L1 in the HFF from collected from left and right ovaries from 3 unique patients. Whole gel Amidoblack total protein staining matching PD-1 and PD-L1 immunoblots in (a) is included for evaluation of Exosome (E) versus Soluble protein (S) fractions. E fractions contained detectable PD-1 and PD-L1, while S fractions did not contain detectable PD-1 or ligands; representative blot for PD-L2 is shown in panel b. Molecular weight standards are marked with black bars and labeled.

We next addressed the question of whether HFF can exhibit physiological immunomodulatory activity, and whether this relates to soluble PD-1 pathway factor concentration(s). For these experiments, we added different volumes of HFF or PBS vehicle to human T cells (isolated separately, in 50 *µ*l culture media) that were optionally activated using a standard anti-CD3/anti-CD28 regime (CD3/CD28 activated)^49^. Interferon gamma (IFN*γ*) production and tyrosine 248 phosphorylation of PD-1 (P-Y248 PD-1) were monitored as measures of relative T cell activation^50,51^ and ligand interaction with T cell PD-1 receptors, respectively (Fig. 6).

**Figure 6.**
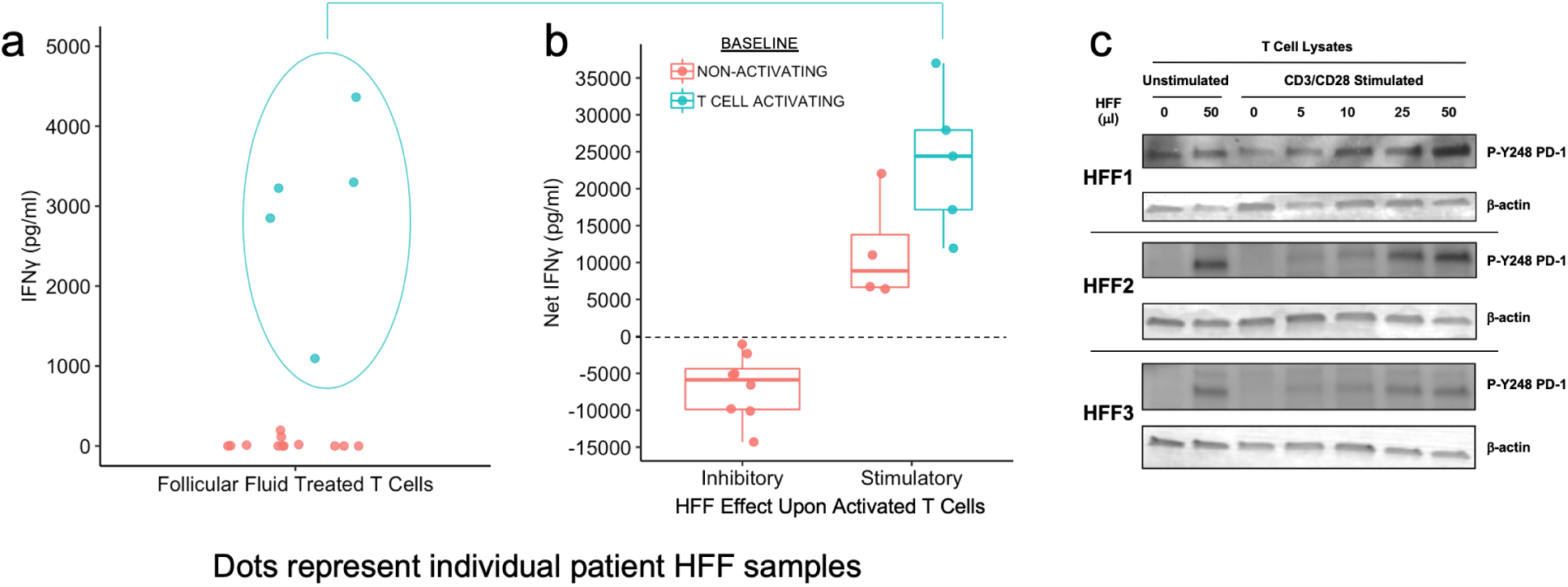
Bioactivity of follicular fluid upon T cell activation. T cells were incubated with hFF without (a) or with (b) concurrent CD3/CD28 activation and IFN*γ* was measured 24 hours later. Critically, IFN*γ* was not detectable in any hFF sample prior to addition to T cells (not shown). In non-activated T cells, some hFF samples stimulated IFN*γ* production above baseline (blue points within oval; mean values for 3 replicates shown). When the same hFF samples were applied to CD3/CD28-activated T cells, net IFN*γ* production was calculated by subtracting the mean amount produced by vehicle-treated, activated controls (set to zero, dashed line). Compared to vehicle, IFN*γ* production was either diminished (“Inhibitory”) or enhanced (“Stimulatory”) by hFF treatment. All 5 samples shown to stimulate IFN*γ* production in non-activated T cells (a) were found to have “Stimulatory” action upon activated T cells (b). Panel c shows western blots of T cell lysates after the described treatments, probed for “active” P-Y248 PD-1 and *β*-actin housekeeping protein. hFF treatment corresponded to increased P-Y248 PD-1 in both Unactivated and Activated T cells, with a volume-dependent dose-response apparent in Activated T cells.

The effect of HFF upon T cell IFN*γ* production varied between patients (n=16, Fig. 5). This was first seen in samples where “naïve” T cells were treated with HFF but were not activated by the anti-CD3/anti-CD28 regime (5A). While all of these T cell samples treated with PBS vehicle did not produce detectable IFN*γ* (not shown), five of sixteen samples treated with HFF (5/16) produced detectable IFN*γ* above background (2162.9 ± 635.1 pg/ml; mean ± SEM). Next, we assessed T cells that were CD3/CD28-activated. To determine the net effect of HFF treatment upon IFN*γ*, we subtracted the mean IFN*γ* produced from activated T cells treated with PBS vehicle from the amount produced from T cells treated with the individual HFF samples. This calculation revealed that HFF could either induce a net increase or decrease in IFN*γ* relative to that control (Fig. 5B). Of the 16 samples, 9 were found to be “stimulatory” and 7 were found to be “inhibitory” of IFN*γ* production. It was notable that all 5 of the samples that induced IFN*γ* in unstimulated T cells (5A) were “stimulatory”, and enhanced the effect of CD3/CD28 activation upon IFN*γ* production (5B).

Last, we addressed the question of whether T cell PD-1 signaling was modulated by HFF addition. Here we measured P-Y248 PD-1 in T cells treated with varying amounts of HFF. Western blots of T cell lysates (unstimulated or CD3/CD28-stimulated) collected after treatment with PBS vehicle of HFF are shown for three different patient HFF samples. HFF was shown to increase P-Y248 PD-1 in both unstimulated and stimulated T cells. Further, there was a correspondence between increasing HFF volume and P-Y248 PD-1 in CD3/CD28-stimulated T cells. We interpret these data as showing that soluble ligands are indeed present at bioactive levels, at least in this PD-1 phosphorylation assay. Given the consistent T cell P-Y248 PD-1 response to HFF treatment, we expect that the cytokine profile of individual HFF specimens is critical in determining the ultimate T cell and overall immune response in real time, and this is being explored in ongoing studies.

## Discussion

We show herein that membrane bound and soluble PD-1 pathway receptor and ligand proteins are expressed in resident (non-lymphoid/non-myeloid) cells of the human and mouse ovary, and can be found in the fluid collected from peri-ovulatory follicles during IVF retrieval procedures. Full-length PD-1 and ligands were found to be highly enriched in exosome fractions in HFF. We further show that these factors (particularly PD-L1) are present in variable amounts within human ovarian cortex pre- and postmenopausally, in a pattern that is consistent with the increased incidence of EOC after menopause. Why the distribution of PD-L1 is more extensive within the ovarian cortex (e.g., nearer the surface) after menopause is currently unknown. Finally, we also show that the soluble ligands are present in HFF at “bioactive” levels capable of regulating T cell activation, perhaps impacting immune surveillance at the time of ovulation *in vivo*.

Our data evaluating net T cell activation in the context of different concentrations of soluble PD-1 pathway members evoke recent findings of Karunarathne et al.^52^. That group showed that PD-L1 and PD-L2 have *opposing* actions upon T cell activation in a (mouse) model of malaria (see^53^ for a review). They showed that as expected, PD-L1:PD-1 interaction suppresses T cell activation, in that case rendering mice susceptible to malarial death. In contrast, PD-L2 and sPD-L2 delivery rescued this lethality, and additional experiments showed that the rescue resulted from the blocking of PD-L1’s interaction with the receptor by PD-L2. sPD-L2 is consistently present at higher concentrations than sPD-L1 in HFF. In combination, these data predict that ovarian immune surveillance is likely to be enhanced by sPD-L2, and that this can be blocked in when sPD-1 production is high. The net impact of HFF upon immune cell action can be appreciated to involve these PD-1 checkpoint factors as well as their cytokine “profiles” (below).

PD-1 pathway regulatory activity in HFF specifically *via* the T cell-activating soluble ligands, and the effect of concurrent soluble PD-1 expression have important implications. First, it has been reported that sPD-1^Δex3^ variant is not produced in “healthy individuals”^8,54–56^. Ovarian granulosa cells can instead be recognized as a contributing, physiological source of soluble receptor, including the full-length protein found to be enriched in exosomes. Second, if pathway members regulate the surveillance and elimination of transformed cells *in vivo*, individual variability in physiological (e.g., ovarian including HFF) levels may contribute to individual variability in ovarian cancer risk (below). This concept has been evaluated in the context of rheumatoid arthritis risk^55^, and PD-L1 has been shown to be a prognostic factor in established ovarian cancer patients^57^. Interestingly, sPD-L1 can be generated by proteolytic cleavage^21,22^ as well as alternative splicing, and we will need to determine how and under what circumstances spliced variants are produced in ovarian follicles.

Immune cells have been isolated from HFF samples^58^, and they may contribute soluble PD-1 and ligands to the fluid prior to and at the time of collection. While we confirmed that granulosa cells can express soluble PD-1 pathway proteins (Fig. 4i-k) using multiple approaches, the relative amount of these proteins that originate from immune cells versus non-immune ovarian cells (e.g., *theca*, granulosa, oocyte) remains unclear. Despite immune cells being found outside of intact, growing ovarian follicles, they may contribute to the protein content of HFF *in situ*, and are likely to enter the HFF sample when the wall of the follicle and its neighboring blood and lymph vessels are mechanically ruptured during the egg retrieval procedure (see below). In addition, clinical gonadotrophic stimulation has been shown to alter levels of detectable cytokines in follicular fluid and also the complement of immune cells retrieved^59^. This stimulation may also be impacting the expression and action of the PD-1 checkpoint in immune and non-immune cells of the ovary.

The degree to which EOC arises from transformed FTE cells that then engraft at healing ovulation sites^39,40,60^ or initiates from ovary-resident cells^61^ is difficult to determine. Regardless, immune checkpoint regulation of T cell activation by FTE and stroma, or, by cells of the ovary could represent a physiological immune-regulatory mechanism. Ovulatory follicular fluid containing soluble PD-1 pathway members may regulate immune surveillance at the ovarian surface and adjacent extra-ovarian peritoneal space. Suppressed PD-1 signaling at this time^52^ could represent a pro-surveillance, “protective” state. Enhanced PD-1 signaling at the time of ovulation would instead reduce T cell surveillance, potentially enhancing the risk of EOC cell survival. Intriguingly, the significant correlations between soluble PD-1 pathway protein concentrations from the largest follicle of patient left and right ovaries (Fig. S4) suggests that sPD-L1 and -L2 production is tightly regulated within individuals, and may relate to overall HGSOC risk. If so, favoring the pro-surveillance state at the time of ovulation may represent a route towards reducing that risk.

Much more mechanistic information about the regulation of immune surveillance within the ovary pre- and post-menopause (and indeed, the immune cell repertoire of the organ) is needed. Our finding that the distribution of somatic ovarian cortex cells that express PD-L1 changes between pre- and postmenopausal life may relate to the increased incidence of EOC after menopause^41^, and again, different susceptibility in individuals. It is likely that the global decline in numbers of T cells that results from aging^62^ and also, the onset of menopause^63^, contibutes as well. Because the application of immunotherapy (checkpoint blockade therapy, or, chimeric antigen receptor, or, “CAR T”) to HGSOC is in its infancy, these data will inform treatment design so that basal PD-1 action in native cells of the ovary and fallopian tube is accounted for.

The broader-than-expected action of the pathway in the ovary may also reflect an immune-privilege mechanism that reduces auto-immunity to cells of the follicle, including developing oocytes^64–66^. Several groups, including Fahmi et al.^67^ and Kollman et al.^59^ have evaluated HFF immune cell populations and cytokine levels, and the potential impact of altered ovarian immune function upon the likelihood of assisted conception. Data from the latter paper showed that gonadotrophic stimulation alters the cytokine profile of isolated HFF compared to HFF collected after a “natural cycle,” albeit where the “natural cycle” included a hormonal (hCG) ovulation trigger. Our direct experimental evaluation of immunomodulation of T cell activity by HFF (Fig. 5) suggests that immune checkpoint signaling may impact ovarian function and reproductive outcomes in combination with cytokine expression and response.

Overall, we interpret these findings as suggestive of a PD-1 pathway-regulated balance between protecting the ovary, including its oocytes and embryos passing through the fallopian tube, from autoimmune reactions, while ensuring that the immune system can function in these organs to protect against infection and malignant transformation.

## Materials and Methods

### Tissue, cell, and follicular fluid collection and processing

All methods were carried out in accordance with appropriate guidelines and regulations. Specifically, approval for human sample provision and collection was granted by the Colorado Multiple Institutional Review Board as de-identified, discarded tissue (Johnson COMIRB Protocol #17-1428). Paraffin-embedded human ovary specimens were generated using a standard protocol by the University of Colorado Department of Pathology. Mouse tissues were collected under the auspices of protocols 347 and 569 (approved by CU-Anschutz IACUC, Bitler).

HFF and HCGC samples were isolated at the University of Colorado Center for Advanced Reproductive Medicine during egg retrieval procedures. Aspirates from the largest follicle within each ovary were collected, and removal of the oocyte and proximal-most HCGC was performed by mechanical cutting. Remaining cumulus cells within HFF were then stored at room temperature (RT) until batch processing to separate GC from HFF by centrifugation (5 minutes, 2000 × g). Cell-free HFF was removed from the cell pellet, and was either aliquoted, and frozen, or processed further for exosome preparations (below). Resuspended HCGC were either flash-frozen or placed into culture media.

Cell-free HFF was optionally sub-fractionated into exosome-rich or exosome-depleted (soluble protein) fractions as follows. HFF was processed using a exosome isolation kit (Total Exosome Isolation Kit, Invitrogen, #4484450 and Total Exosome Isolation Reagent, Invitrogen #4484453) per the manufacturer’s instructions.

Ovaries from 8-week-old C57Bl/6 mice were isolated, rinsed in PBS, and placed into 4% paraformaldehyde prior to further processing (*Immunostaining*, below).

### PD-1 pathway gene expression analyses (immunostaining, western blots, and ELISA assays) Immunostaining

Ovary, fallopian tube, or control tonsil samples were embedded in paraffin and sectioned. Clinical specimens provided for research purposes were sectioned by the University of Colorado Pathology Research Core service. Briefly, after deparaffinization and rehydration, antigen retrieval was performed by incubating the sections in 10 mM citrate buffer pH 6 at 120*°*C for 30 min. Endogenous peroxidase blocking was performed by using 3% hydrogen peroxide (Labchem, Cat # LC154301). Sections were then washed in water, then in PBS containing 0.05% Tween 20 (0.05% TBST). Tissue sections were blocked in 1% (wt/vol) bovine serum albumin plus 10% normal goat serum at room temperature (RT) for 90 min. After blocking, the tissues were incubated overnight at 4*°*C with primary antibody in a humid chamber. After three washes 0.05 % TBST, sections were incubated for 30 min goat anti-rabbit (abcam, #6721). Signal amplification was performed using an ABC Kit (Vectastain, #PK6100) for 30 min at RT followed by DAB (Vector Labs, #SK-4100) substrate. Nuclei were counterstained with Harris hematoxylin for 2 min. Following rinsing, slides were dehydrated and mounted in Permafluor (Fisher Scientific, # SP15-500). Negative control sections were submitted to the same procedures, except that the first antibody was replaced by blocking solution. Pictures were taken using an Olympus microscope (Model: CKX41SF, SN:2B77130)

### Western Blot Analyses

Protein isolation from whole ovaries was performed by using TPER Tissue Protein Extraction (Thermo Scientific, #78510). Protease Inhibitor Cocktail (Sigma, #P8340) and Phosphatase Inhibitor Cocktail (Calbiochem, #524625) were each added at a 1:100 dilution. Where applicable, immunoprecipitation was performed using lysis buffer (Cell Signaling Technologies, #9803) and Protein A (#9863) according to the manufacturer’s protocol. Tissue or cell digestion was performed by agitation with stainless steel beads (NextAdvance, #SSB14B-RNA) for 5 min at speed 12 at 4*°*C in a bullet blender. Supernatant was recovered and the Bradford assay was performed for protein quantification. Supernatants were electrophoresed on a 4–12% polyacrylamide gel (Invitrogen), and proteins were transferred to a HyBond PVDF membrane (Amersham). Membranes were blotted with antibodies against the protein targets in the summary table below. HRP-conjugated second antibody was used followed by chemi-luminescence imaging using X-ray film. Optionally, a near-IR second antibody (LI-COR) was used and blots were imaged using the LI-COR Odyssey imaging system and densitometric analysis was performed using ImageStudio (LI-COR). For experiments comparing exosome and soluble proteins in fractions of the same HFF specimen (HFF processing, above), equal concentrations of exosome and soluble protein fractions from each sample were loaded onto SDS-page gels so that the relative amount of protein in each fraction could be estimated.

### ANTIBODIES / SUPPLIERS

| Target | Supplier | Catalog # |
| --- | --- | --- |
| **HUMAN** |  |  |
| PD-1 | Cell Signaling Technology | #86163 |
| PD-1<sup>†</sup> | R&D Systems | #AF1086 |
| PD1 (phospho Y248) | abcam | #ab206378 |
| PD-L1 | ThermoFisher | #PA5-20343 |
| PD-L2 | BIO-RAD | AHP1704 |
| **MOUSE** |  |  |
| Pd-1 | ProSci, Inc. | #4065 |
| Pd-L1<sup>‡</sup> | ThermoFisher | #PA5-20343 |
| Pd-L2<sup>‡</sup> | BIO-RAD | #AHP1704 |

† - Blocking antibody for T cell assays
‡ - Same antibody cross-reacts with human and mouse.

### ELISA Assays

ELISA assays were performed according to manufacturer’s instructions. Each sample was run in triplicate, and each assay included positive and negative controls in addition to standard curve and “unknown” HFF samples.

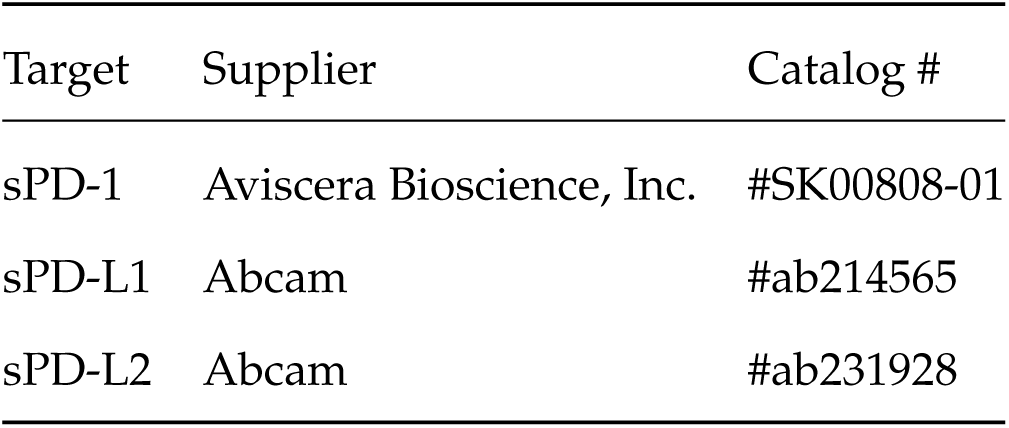

### T cell isolation, handling, and activation assay

T cell provision and analyses were performed by the Human Immune Monitoring Shared Resource at CU|AMC. The source of T cells was from a Bonfils Leukopak (LRS chamber), first sorted with a Miltenyi pan T cell isolation kit, and then sorted on the AutoMacs instrument. Media for cell culture and activation or control treatment were RPMI + 10% FBS + 2 mM + L-glut + 200U/ml penicillin/streptomycin. T cell activation was performed by anti-CD3/anti-CD28 treatment (Supplier, #s). Where specified, PD-1 blocking antibody (R&D Systems/Techne, #AF1086) was included at the 10 *µ*g/ml. IFN*γ* levels were measured using the MSD V-PLEX assay (Mesoscale, Inc., # K151QOD) according to manufacturer’s instructions. HFF samples were diluted 50-fold in assay kit Diluent 2 for the assay.

## Acknowledgements

Drs. Jill Slansky, Kian Behbahkt, Saketh Guntupalli, and Bradley Corr are acknowledged for their suggestions while the manuscript was being developed. Melissa Rosario and David Russell collected human follicular fluid and granulosa-lutein cells discarded after clinical IVF procedures and are gratefully acknowleged for their efforts. Jennifer Ann McWilliams and Angela Minic performed T cell activation assays. Dr. Sunhyo Ryo performed “active” P-Y248 T cell western blots.

## Author Contribution Statement

J.J. and B.G.B wrote the main manuscript text, and J.J. performed statistical analyses; P.K.S., R.A., E.L.C., E.S.B., T-H.L., L.P., and K.J. performed experiments and processed original data. A.N.K., L.N-T., A.J.P., and M.D.P. performed study design for clinical specimens; M.D.P. and D.J.O. provided histopathological assessment of study slides.

