## Supplemental Figures for "Expression and T cell Regulatory Action of the PD-1 Immune Checkpoint in the Ovary and Fallopian Tube"

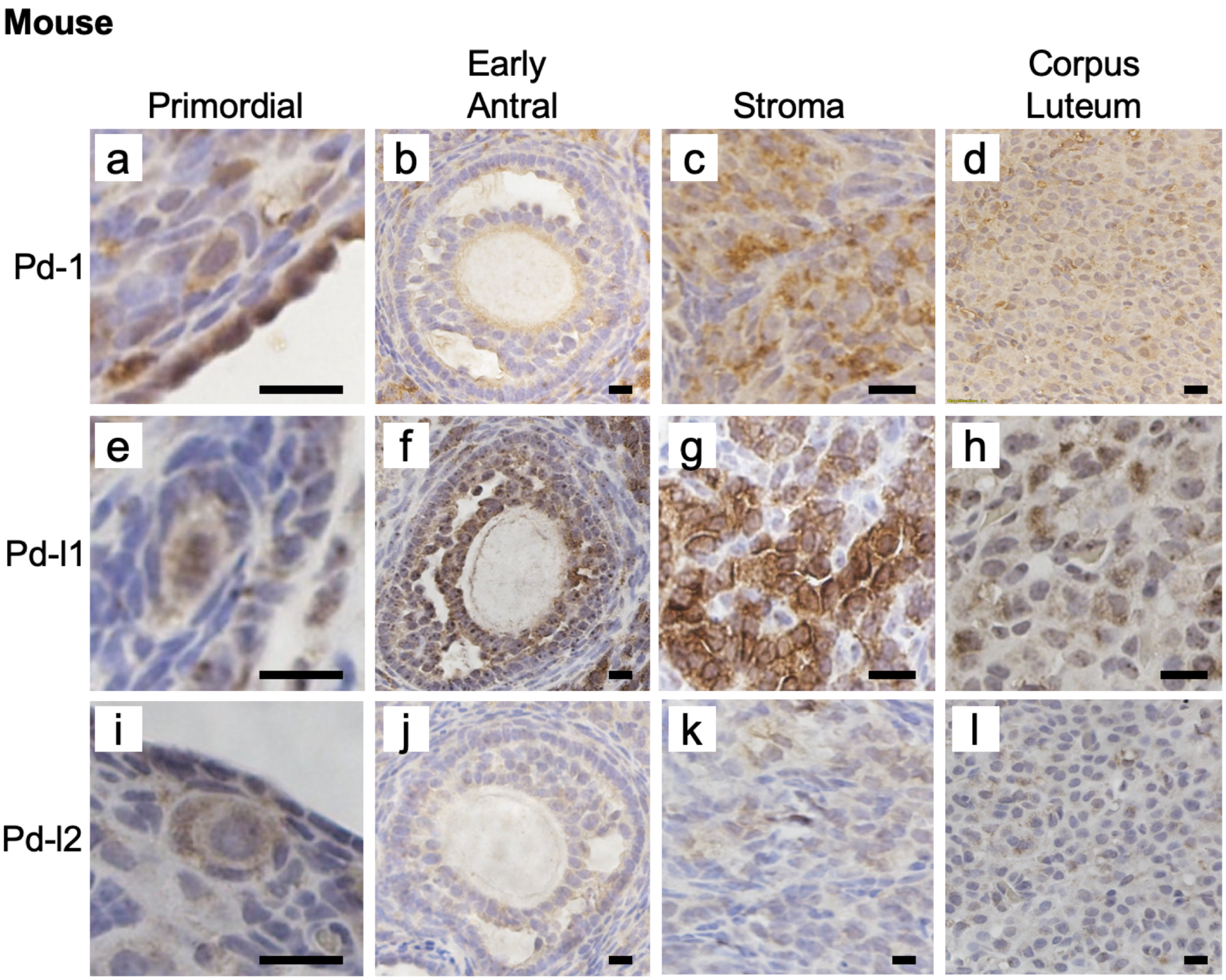


**Figure S1. Expression of Pd-1 pathway proteins in non-lymphoid and non-myeloid cells of the adult mouse ovary**. Colorimetric immunostaining was performed on n=3 unique adult (9-12 week-old) mouse ovary specimens, brown is positive signal. PD-1 (a-d), PD-L1 (e-h), and PD-L2 (i-l) were consistently and broadly expressed in cells of ovarian follicles, ovarian stroma, and *corpora lutea*. Scale bars = 10 μm.

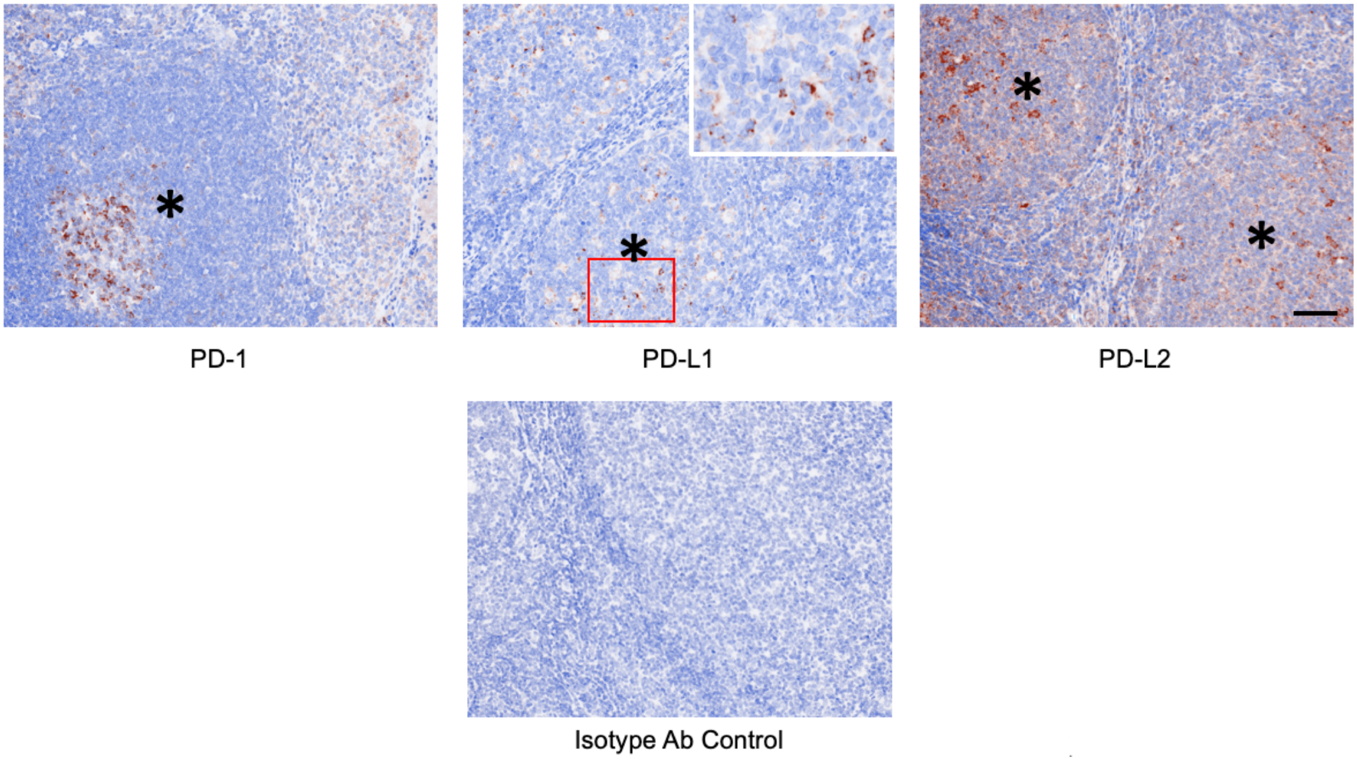


**Figure S2. Control immunostaining of human tonsil tissue for PD-1 receptor and ligands.** Primary antibodies used for ovary and fallopian tube staining were also used to probe human tonsil tissue as a positive control. Cells positive for PD-1, PD-L1, and PD-L2 were detected in these specimens (brown staining). Tonsil germinal centers are marked by asterisks (*). Inset in PD-L1 panel is region bounded by red box. Scale bar in PD-L2 panel = 150 µm applies to all images.

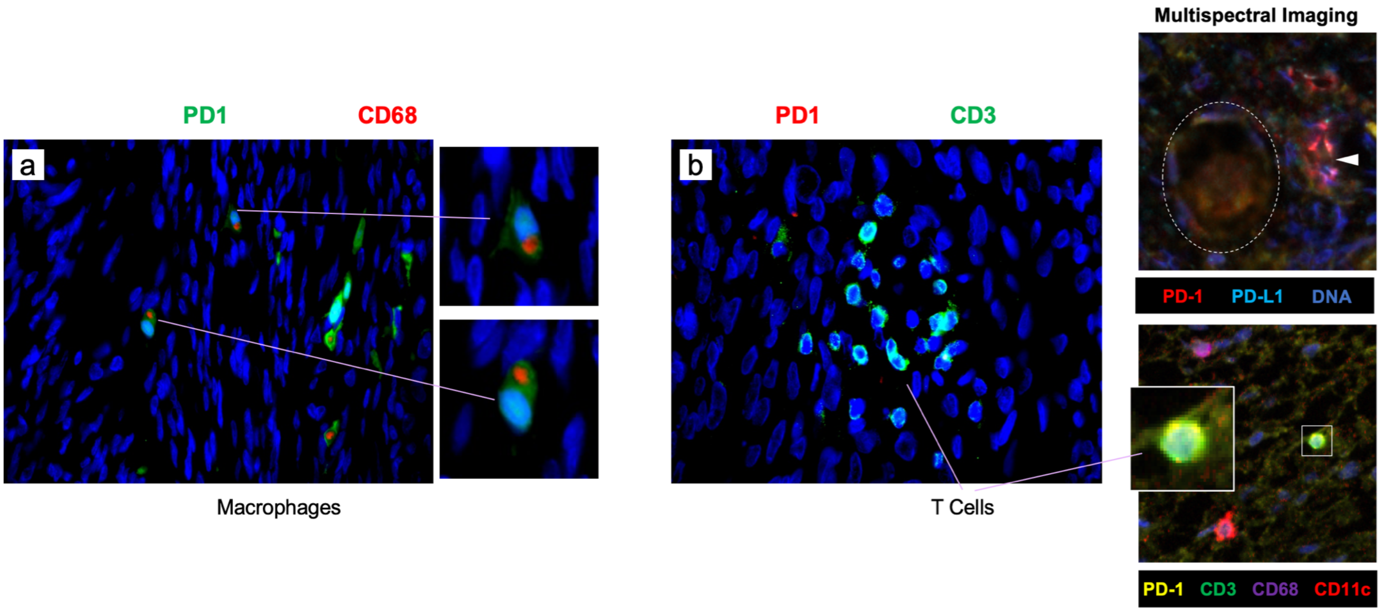


**Figure S3. Multichannel labeling analysis of PD-1-positive cells and immune cell markers**. Small, intensely PD-1 positive cells (colorimetric staining, insets, Fig. 1b,d; 2a) in the human ovary were hypothesized to be immune cells. Panel a shows a representative image from dual staining of n=3 unique ovary specimens was performed for PD-1 (green) and CD68 (red, left panels, individual cells shown at higher magnification). Double-labeled cells, with intracellular CD68 “patches” reminiscent of the reported Golgi apparatus and vesicle distribution in macrophages (https://www.proteinatlas.org/ENSG00000129226-CD68/cell#human) were identified in all specimens. T cell staining using anti-CD3 (panel b) revealed individual (not shown) and "patches" of T cells in the ovarian cortex and medulla, and T cells were mostly negative for PD-1. Vectra multispectral imaging of the human ovary (c) was also performed using different primary antibodies to PD-1 pathway proteins. We confirmed PD-1 and PD-L1 expression in small follicles (top image, follicle within dashed oval) with simultaneous imaging of PD-1-positive non-follicular cells (arrowhead). In addition to rare PD-1 positive T cells (CD3 – green, PD-1 – yellow; see electronically magnified inset), we were able to simultaneously detect dendritic cells (CD11c – red) and macrophages (CD68, purple).

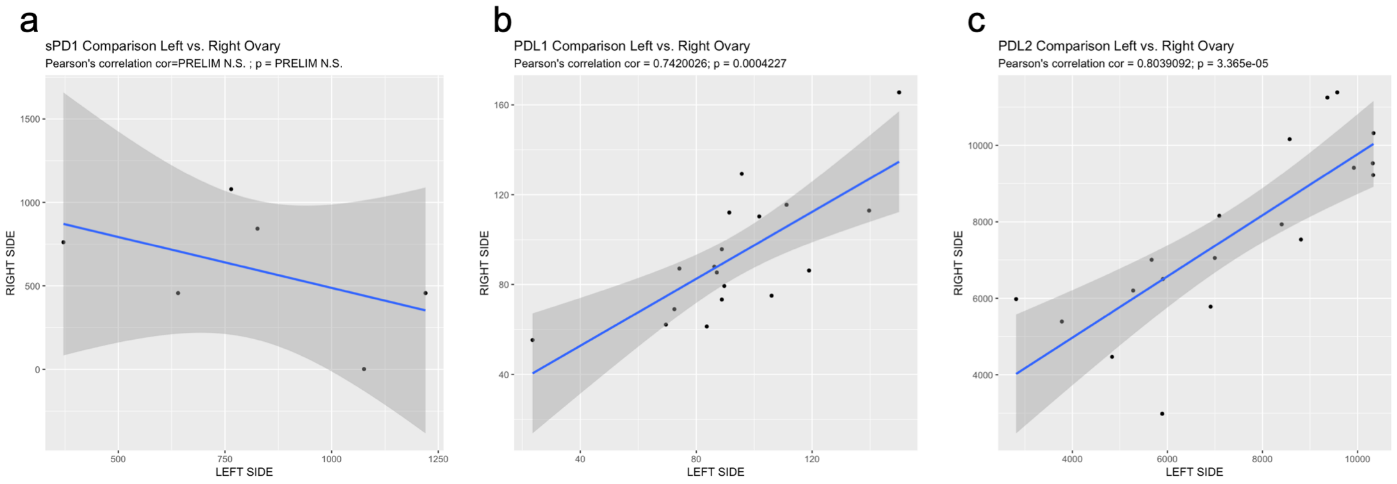


**Figure S4. Correlation analysis of PD-1 and ligands between samples from left and right ovary.** Levels (pg/ml) of PD-1 (a), PD-L1 (b), and PD-L2 (c) from HFF samples where the largest follicle from left and right ovary were collected were determined (see also Fig. 4a-c, Fig. S5). Correlation analysis was performed between the sides, and Pearson's correlation coefficient was calculated for all data points and is shown in the subtitle of each plot along with the calculated p value.

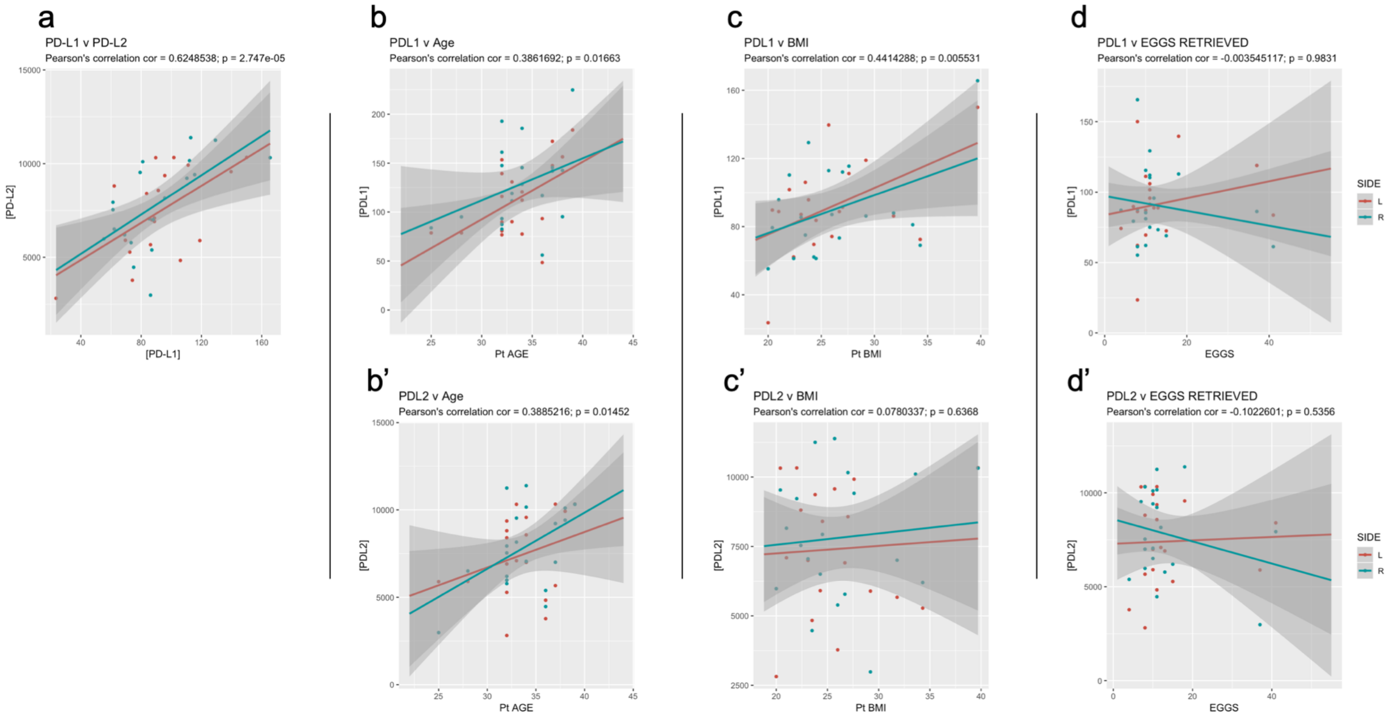


**Figure S5. Further correlation analysis of PD-1 ligands.** Levels (pg/ml) of PD-L1 and PD-L2 from HFF samples where the largest follicle from left (L) and right (R) ovary were collected were compared to each other (panel a, PD-L1 v PD-L2), and compared to patient age (b), and characteristic (c - body mass index [BMI], d - number of eggs retrieved) data. Pearson's correlation coefficient was calculated for all data points (calculated from combined L and R side data) and is shown in the subtitle of each plot. Data points and correlation lines are shown separately for L side HFF (red) and R side HFF (blue). Significant correlations (Pearson's correlation test) were detected between the concentrations of PD-L1 and PD-L2 (a), between both ligands and patient age (b, b'), and between PD-L1 and patient BMI ($p < 0.05$).
